# Adaptive social contact rates induce complex dynamics during epidemics

**DOI:** 10.1101/2020.04.14.028407

**Authors:** Ronan F. Arthur, James H. Jones, Matthew H. Bonds, Yoav Ram, Marcus W. Feldman

**Affiliations:** School of Medicine, Stanford University, Stanford, CA, USA; Department of Earth Systems Science, Stanford University, Stanford, CA, USA; Department of Global Health and Social Medicine, Harvard Medical School, Cambridge, MA, USA; Efi Arazi School of Computer Science, IDC Herzliya, Herzliya, Israel; Department of Biology, Stanford University, Stanford, CA

## Abstract

The COVID-19 pandemic has posed a significant dilemma for governments across the globe. The public health consequences of inaction are catastrophic; but the economic consequences of drastic action are likewise catastrophic. Governments must therefore strike a balance in the face of these trade-offs. But with critical uncertainty about how to find such a balance, they are forced to experiment with their interventions and await the results of their experimentation. Models have proved inaccurate because behavioral response patterns are either not factored in or are hard to predict. One crucial behavioral response in a pandemic is adaptive social contact: potentially infectious contact between people is deliberately reduced either individually or by fiat; and this must be balanced against the economic cost of having fewer people in contact and therefore active in the labor force. We develop a model for adaptive optimal control of the effective social contact rate within a Susceptible-Infectious-Susceptible (SIS) epidemic model using a dynamic utility function with delayed information. This utility function trades off the population-wide contact rate with the expected cost and risk of increasing infections. Our analytical and computational analysis of this simple discrete-time deterministic model reveals the existence of a non-zero equilibrium, oscillatory dynamics around this equilibrium under some parametric conditions, and complex dynamic regimes that shift under small parameter perturbations. These results support the supposition that infectious disease dynamics under adaptive behavior-change may have an indifference point, may produce oscillatory dynamics without other forcing, and constitute complex adaptive systems with associated dynamics. Implications for COVID-19 include an expectation of fluctuations, for a considerable time, around a quasi-equilibrium that balances public health and economic priorities, that shows multiple peaks and surges in some scenarios, and that implies a high degree of uncertainty in mathematical projections.

**Author summary:** Epidemic response in the form of social contact reduction, such as has been utilized during the ongoing COVID-19 pandemic, presents inherent tradeoffs between the economic costs of reducing social contacts and the public health costs of neglecting to do so. Such tradeoffs introduce an interactive, iterative mechanism which adds complexity to an infectious disease system. Consequently, infectious disease modeling typically has not included dynamic behavior change that must address such a tradeoff. Here, we develop a theoretical model that introduces lost or gained economic and public health utility through the adjustment of social contact rates with delayed information. We find this model produces an equilibrium, a point of indifference where the tradeoff is neutral, and at which a disease will be endemic for a long period of time. Under small perturbations, this model exhibits complex dynamic regimes, including oscillatory behavior, runaway exponential growth, and eradication. These dynamics suggest that for epidemic response that relies on social contact reduction, secondary waves and surges with accompanied business re-closures and shutdowns may be expected, and that accurate projection under such circumstances is unlikely.

## Introduction

The COVID-19 pandemic had infected almost 9 million people and caused over 450,000 deaths worldwide as of June 23, 2020 [1]. In the absence of effective therapies and vaccines [2], many governments responded with lock-down policies and social distancing laws to reduce the rate of social contacts and curb transmission of the virus. Prevalence of COVID-19 in the wake of these policies in the United States indicates they may have been successful at decreasing the reproduction number (*R*_*t*_) of the epidemic [1].

However, they have also led to economic recession with an unemployment rate at an 80-year peak, the stock market in decline, and the federal government forced to borrow heavily to financially support businesses and households. Solutions to these economic crises may conflict with public health recommendations. Thus, governments worldwide must decide how to balance the economic and public health consequences of their epidemic response interventions.

Behavior-change in response to an epidemic, whether autonomously adopted by individuals or externally directed by governments, affects the dynamics of infectious diseases [3, 4]. Prominent examples of behavior-change in response to infectious disease prevalence include measles-mumps-rubella (MMR) vaccination choices [5], social distancing in influenza outbreaks [6], condom purchases in HIV-affected communities [7], and social distancing during the ongoing COVID-19 pandemic [2]. Behavior is endogenous to an infectious disease system because it is, in part, a consequence of the prevalence of the disease, which in turn responds to changes in behavior [8, 9].

Individuals and governments have greater incentive to change behavior as prevalence increases; conversely they have reduced incentive as prevalence decreases [10, 11]. Endogenous behavioral response may then theoretically produce a non-zero endemic equilibrium of infection. This happens because, at low levels of prevalence, the cost of avoidance of a disease may be higher than the private benefit to the individual, even though the collective, public benefit in the long-term may be greater. However, in epidemic response we typically think of behavior-change as an exogenously-induced intervention without considering associated costs. While guiding positive change is an important intervention, neglecting to recognize the endogeneity of behavior can lead to a misunderstanding of incentives and a resurgence of the epidemic when behavior change is reversed prematurely.

Although there is growing interest in the role of adaptive human behavior in infectious disease dynamics, there is still a lack of general understanding of the most important properties of such systems [3, 8, 12]. Behavior is difficult to measure, quantify, or predict [8], in part, due to the complexity and diversity of human beings who make different choices under different circumstances. Despite these challenges, modelers and mathematicians have adopted a variety of strategies to investigate the importance of behavior (see Funk et al., 2010 [4] for a general review). For example, an early expansion of the Kermack-McKendrick susceptible-infectious-removed (SIR) model simply allowed the transmission parameter (*β*) to be a negative function of the number infected, effectively introducing an intrinsic negative feedback to the infected class that regulated the disease [13].

Modelers have used a variety of tools, including agent-based modeling [14], network structures for the replacement of central nodes when sick [15] or for behavior-change as a social contagion process [16], game theoretic descriptions of rational choice under changing incentives as with vaccination [6, 11, 17], and a branching process for heterogeneous agents and the effect of behavior during the West Africa Ebola epidemic in 2014 [18]. A common approach to incorporating behavior into epidemic models is to track co-evolving dynamics of behavior and infection [16, 19, 20], where behavior represents an *i*-state of the model [21]. In a compartmental model, this could mean separate compartments (and transitions therefrom) for susceptible individuals in a state of fear and those not in a state of fear [16].

Periodicity (i.e. multi-peak dynamics) has long been documented empirically in epidemiology [22, 23]. Periodicity can be driven by seasonal contact rate changes (e.g. when children are in school) [24], seasonality in the climate or ecology [25], sexual behavior change [26], and host immunity cycling through new births of susceptibles or a decay of immunity over time. Some papers in nonlinear dynamics have studied delay differential equations in the context of epidemic dynamics and found periodic solutions [27]. Although it is atypical to include delay in modeling, delay is an important feature of epidemics. For example, if behavior responds to mortality rates, there will inevitably be a lag with an average duration of the incubation period plus the life expectancy upon becoming infected. In a tightly interdependent system, reacting to outdated information can result in an irrational response and periodic cycling.

A delay in a behavioral response to an epidemic can take a variety of forms. In the ongoing COVID-19 pandemic, for example, there have been delays in the international recognition of the outbreak, delays in the identification of the virus, delays in the acquisition of reliable information on suspected and confirmed cases, delays in the development and deployment of testing, delays in the launching of community information and communication campaigns, and delays in laying the physical infrastructure for the treatment and isolation of an increasing number of patients. Health authorities have had to wait up to 14 days to see the epidemiological consequences of public gatherings, opening of businesses, and guidelines and policies regarding personal protective equipment. Recognizing the effects of such delays as a part of the endogenous relationship between disease and behavior is important to improved understanding of the complexity of disease dynamics.

The original epidemic model of Kermack and McKendrick [28] was first expressed in discrete time. Then by allowing “the subdivisions of time to increase in number so that each interval becomes very small” the famous differential equations of the SIR epidemic model were derived. Here we begin with a discrete-time Susceptible-Infected-Susceptible model that is adjusted on the principle of endogenous behavior-change through an adaptive social-contact rate that can be thought of as either individually motivated or institutionally imposed. We introduce a dynamic utility function that motivates the population’s effective contact rate at a particular time period. This utility function is based on information about the epidemic size that may not be current. This leads to a time delay in the contact function that increases the complexity of the population dynamics of the infection. Results from the discrete-time model show that the system approaches an equilibrium in many cases, although small parameter perturbations can lead the dynamics to enter qualitatively distinct regimes. The analogous continuous-time model retains periodicities for some sets of parameters, but numerical investigation shows that the continuous time version is much better behaved than the discrete-time model. This dynamical behavior is similar to models of ecological population dynamics, and a useful mathematical parallel is drawn between these systems.

## Model specifications

### SIS

To represent endogenous behavior-change, we start with the classical discrete-time susceptible-infected-susceptible (SIS) model [28], which, when incidence is relatively small compared to the total population [29, 30], can be written in terms of the recursions

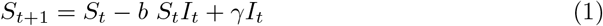

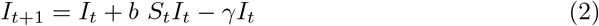

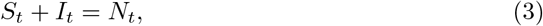

where at time t, *S*_*t*_ represents the number of susceptible individuals, *I*_*t*_ the infected individuals, and *N*_*t*_ the number of individuals that make up the population, which is assumed fixed in a closed population. We can therefore write N for the constant population size. Here *γ*, with 0 < *γ* < 1, is the rate of removal from *I* to *S* due to recovery. This model in its simplest form assumes random mixing, where the parameter *b* represents a composite of the average contact rate and the disease-specific transmissibility given a contact event. In order to introduce human behavior, we substitute for *b* a time-dependent *b*_*t*_, which is a function of both *b*_0_, the probability that disease transmission takes place on contact, and a dynamic social rate of contact *c*_*t*_ whose optimal value, 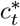, is determined at each time *t* as in economic epidemiological models [31], namely

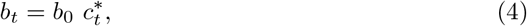

where 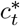 represents the optimal contact rate, defined as the number of contacts per unit time that maximize utility for the individual. Here, 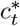 is a function of the number of infected in the population according to the perceived risks and benefits of social contacts, which we model as a utility function, assuming there is a constant utility independent of contact, a utility loss associated with infection, and a utility derived from the choice of number of daily contacts with a penalty for deviating from the choice of contacts which would yield the most utility.

This utility function is assumed to take the form

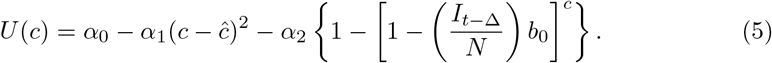

Here *U* represents utility for an individual at time t given a particular number of contacts per unit time *c*, *α*_0_ is a constant that represents maximum potential utility achieved at a target contact rate *ĉ*. The second term, *α*_1_(*c* − *ĉ*)^2^, is a concave function that represents the penalty for deviating from *ĉ*. The third term, 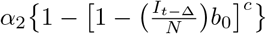, is the cost of infection (i.e. morbidity), *α*_2_, multiplied by the probability of infection over the course of the time unit. The time-delay Δ represents the delay in information acquisition and the speed of response to that information. We note that 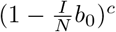 can be approximated by

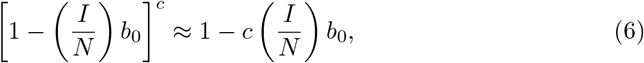

when 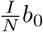 is small and 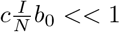. We thus assume 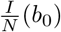 is small, and approximate *U* (*c*) in Eq. 5 using Eq. 6. Eq. 5 assumes a strictly negative relationship between number of infecteds and contact.

We assume an individual or government will balance the cost of infection, the probability of infection, and the cost of deviating from the target contact rate *ĉ* to select an optimal contact rate 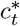, namely the number of contacts, which takes into account the risk of infection and the penalty for deviating from the target contact rate. This captures the idea that individuals trade off how many people they want to interact with versus their risk of getting sick, or that authorities want to reopen the economy during a pandemic and have to trade off morbidity and mortality from increasing infections with the need to allow additional social contacts to help the economy restart. This optimal contact rate can be calculated by finding the maximum of *U* with respect to *c* from Eq. 5 with substitution from Eq. 6, namely

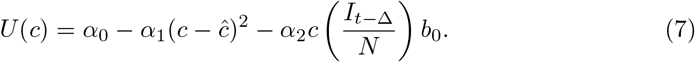

Differentiating, we have

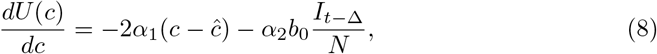

which vanishes at the optimal contact rate, *c*\*, which we write as 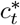 to show its dependence on time. Then

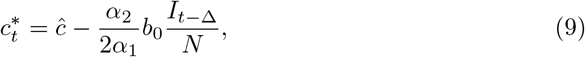

which we assume to be positive. Therefore, total utility will decrease as *I*_*t*_ increases and 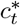 also decreases. Utility is maximized at each time step, rather than over the course of lifetime expectations. In addition, Eq 9 assumes a strictly negative relationship between number of infecteds at time *t* − Δ and *c*\*. While behavior at high degrees of prevalence has been shown to be non-linear and fatalistic [32, 33], in this model, prevalence (i.e., 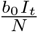) is assumed to be small, consistent with Eq. 6.

We introduce the new parameter 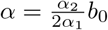, so that

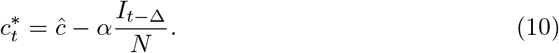

We can now rewrite the recursion from Eq. 2, using Eq. 4 and replacing *c*_*t*_ with 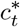 as defined by Eq. 10, as

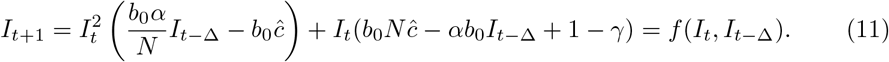

When Δ = 0 and there is no time delay, *f*(·) is a cubic polynomial, given by

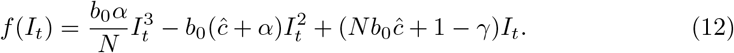

### SIR

For the susceptible-infected-removed (SIR) version of the model, we include the removed category and write the (discrete-time) recursion system as

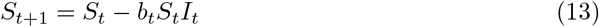

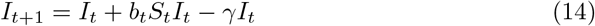

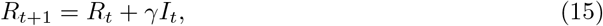

where *R*_*t*_ = *N* − *I*_*t*_ − *S*_*t*_, 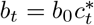 with *b*_0_ the baseline contact rate and 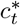 specified by Eq. 10. With *b*_*t*_ = *b*, say, and not changing over time, Eqs. 13–15 form the discrete-time version of the classical Kermack-McKendrick SIR model [28]. The inclusion of the removed category entails that *Ĩ* = 0 is the only equilibrium of the system Eqs. 13–15; unlike the SIS model, there is no equilibrium with infecteds present. In general, since 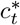 includes the delay Δ, the dynamic approach to *Ĩ* = 0 is expected to be quite complex. Intuitively, since the infecteds are ultimately removed, we do expect that from any initial frequency *I*_0_ of infecteds all *N* individuals will eventually be in the *R* category. Numerical analysis of this SIR model shows strong similarity between the SIS and SIR models for several hundred time steps before the SIR model converges to *Ĩ* = 0 with *R* = *N*. In the section “Numerical Iteration and Continuous-Time Analog” we compare the numerical iteration of the SIS (Eq. 11) and SIR (Eqs. 13–15) and integration of the continuous-time (differential equation) versions of the SIS and SIR models.

## Analytical Results

### Equilibria

To determine the dynamic trajectories of (11) without time delay, we first solve for the fixed point(s) of the recursion (11) (i.e., value or values of *I* such that *f*(*I*_*t*+1_) = *I*_*t*_ = *I*_*t*−Δ_). That is, we solve

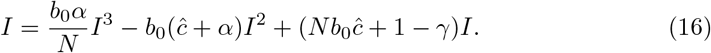

From Eq. 16, it is clear that *I* = 0 is an equilibrium as no new infections can occur in the next time-step if none exist in the current one. This is the disease-free equilibrium denoted by *Ĩ*. Other equilibria are the solutions of

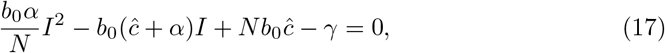

namely

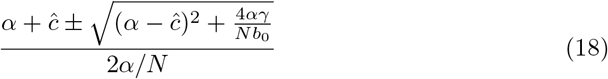

We label the solution with the + sign *I*\* and the one with the − sign *Î*. *I*\* > 0 but *I*\* ≤ *N* if 4*αγ*/*Nb*_0_ ≤ 0, which is impossible under our assumptions that *α* and *γ* are positive. Hence *I*\* is illegitimate. Further, under these same conditions, *Î* ≤ *N*, and *Î* > 0 if

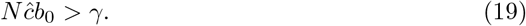

It is important to note that under these conditions *Î* is an equilibrium of the recursion (11) for any Δ ≥ 0. Recall that for the SIR version of this model the only equilibrium is *Ĩ* = 0.

### Stability of the equilibria

Assessing global asymptotic stability in epidemic models is an important task of mathematical epidemiology [34, 35]. The three equilibria of the SIS recursion (11) are qualitatively different. *Ĩ* = 0 corresponds to a disease-free population; *I*\* is greater than *N* and is therefore illegitimate; *Î* is the only positive legitimate equilibrium if *ĉb*_0_ > *γ*/*N* and is, therefore, the most interesting for the asymptotic stability behavior of the epidemic. Mathematical stability analysis of recursion (11) is complicated because of the delay term Δ. However, from (11), if *N*ĉ*b*_0_ > *γ*, the disease-free equilibrium *Ĩ* = 0 is locally unstable, and in this case *Î* is indeed legitimate.

Local stability of *Î* in (18) is discussed in detail in the Appendix. First, in the absence of delay (i.e., Δ = 0), *Î* is locally stable if 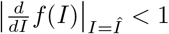, and the condition for this to hold when *Î* is legitimate is

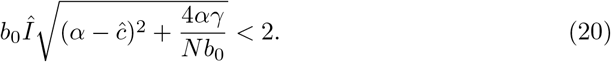

If inequalities (20) and *N*ĉ*b*_0_ > *γ* hold, then *Î* is locally stable. However, even if both of these inequalities hold, the number of infecteds may not converge to *Î*. It is well known that iterations of discrete-time recursive relations, of which (12) is an example (i.e., with Δ = 0), may produce cycles or chaos depending on the parameters and the starting frequency *I*_0_ of infecteds.

Table 1 shows an array of possible asymptotic dynamics with Δ = 0 found by numerical iteration of (12) for a specific set of parameters and an initial frequency *I*_0_ = 1 infected individual. Included in Table 1 are examples for which, beginning with a single infected, the number of infecteds explodes, becoming unbounded; of course, this is an illegitimate trajectory since *I*_*t*_ cannot exceed *N*. However, in the case marked *, *Î* is locally stable and with a large enough initial number of infecteds, there is damped oscillatory convergence to *Î*. In the case marked **, with *I*_0_ = 1 the number of infecteds becomes unbounded, but in this case, *Î* is locally unstable, and starting with *I*_0_ close to *Î* a stable two-point cycle is approached; in this case *df*(*I*)/*dI*|_*I*=*Î*_ < −1.

**Table 1.** Some results for dynamics of infection with Δ = 0.

| Parameters |  |  |  |  | Equilibrium | Dynamics |
| --- | --- | --- | --- | --- | --- | --- |
| $N$ | $b_0$ | $\gamma$ | $\hat{c}$ | $\alpha$ | $\hat{I}$ | |
| 250 | 0.1 | 0.1 | 0.2 | 0.1 | 240.371 | $I_0 = 1$: two-point cycle 110.436, 339.564 |
| 250 | 0.1 | 0.1 | 0.205 | 0.1 | 240.799 | $I_0 = 1$: four-point cycle above and below $\hat{I}$ |
| 250 | 0.1 | 0.1 | 0.209 | 0.1 | 241.115 | $I_0 = 1$: apparent chaos around $\hat{I}$ |
| 250 | 0.5 | 0.1 | 0.1 | 0.1 | 227.639 | $I_0 = 1$: becomes unbounded.<br>$I_0 = 226$: converges to two-point cycle.** |
| 250 | 0.115 | 0.1 | 0.1 | 0.1 | 203.375 | $I_0 = 1$: overshoots $\hat{I}$, then decreases to $\hat{I}$ |
| 350 | 0.1 | 0.1 | 0.1 | 0.1 | 290.839 | $I_0 = 1$: overshoots $\hat{I}$, then decreases to $\hat{I}$ |
| 1,000 | 0.1 | 0.1 | 0.1 | 0.1 | 900.000 | $I_0 = 1$: damped oscillation to $\hat{I}$ |
| 1,100 | 0.1 | 0.1 | 0.1 | 0.1 | 995.119 | $I_0 = 1$: $I_t$ becomes unbounded<br>$I_0 = 990$: damped oscillation to $\hat{I}$ * |
| 10,000 | 0.05 | 0.08 | 0.0015 | 0.375 | 35.718 | $I_0 = 1$: monotone convergence to $\hat{I}$ |

\*, \*\* See text immediately preceding Table 1.

Stability analysis of the SIS model is more complicated when Δ ≠ 0, and in the Appendix we outline the procedure for local analysis of the recursion (11) near *Î*. Local stability is sensitive to the delay time Δ as can be seen from the numerical iteration of (11) for the specific set of parameters shown in Table 2. Some analytical details related to Table 2 are in the Appendix.

**Table 2.** The effect of the delay, Δ, on dynamics of infecteds.*

| $\Delta$ | Outcome |
| --- | --- |
| 0 | Monotone convergence to $\hat{I}$ |
| 1 | Damped oscillation to $\hat{I}$ |
| 2 | $\hat{I}$ locally unstable; $I_0 < 72$ bounded oscillation; $I_0 > 73$ unbounded oscillation |
| 3 | $\hat{I}$ locally unstable; collapse $(-\infty)$ |
| 4 | $\hat{I}$ locally unstable; collapse $(-\infty)$ |

\* In all cases, $N = 10,000$ , $b_0 = 0.05$ , $\gamma = 0.08$ , $\hat{c} = 0.0015$ , $\alpha = 0.375$ , $\hat{I} = 35.718$ . $I_0 = 1$ unless stated.

## Numerical Iteration and Continuous-Time Analog

We begin with numerical analysis of the discrete-time SIS recursion (11), which includes the delay parameter Δ. Local stability properties of the equilibrium state *Î*, with 0 < *Î* < *N*, are shown in the Appendix under the assumption *N*ĉ*b*_0_ > *γ*, which also entails that the disease-free equilibrium *Ĩ* = 0 is locally unstable. In the recursion (11), the number of infecteds at time *t* will not, in general, be integers, as they would be in an actual epidemic. Further, the dynamics of *I*_*t*_ under such a recursion can be very sensitive to the starting condition *I*_0_, the size of the time delay Δ, and the parameters: *N*, *b*_0_, *γ*, *ĉ*, and *α*. The local stability of *Î*, namely whether *I*_*t*_ converges to *Î* from a starting number of infecteds close to *Î*, may tell you little about the actual trajectory of *I*_*t*_ from other starting conditions.

Table 1 reports an array of dynamic trajectories for some choices of parameters and, in two cases, an initial number of infecteds other than *I*_0_ = 1. The first three rows show three sets of parameters for which the equilibrium values of *Î* are very similar but the trajectories of *I*_*t*_ are different: a two-point cycle, a four-point cycle, and apparently chaotic cycling above and below *Î*. In all of these cases, *df*(*I*)/*dI*|_*I*=*Î*_ < −1. Clearly the dynamics are sensitive to the target contact rate *ĉ* in these cases. The fourth and eighth rows show that *I*_*t*_ becomes unbounded (tends to +∞) from *I*_0_ = 1, but a two-point cycle is approached if *I*_0_ is close enough to *Î*: *df*(*I*)/*dI*|_*I*=*Î*_ < −1 in this case. For the parameters in the ninth row, if *I*_0_ is close enough to *Î* there is damped oscillation into *Î*: here −1 < *df*(*I*)/*dI*|_*I*=*Î*_ < 0.

The fifth and sixth rows of Table 1 exemplify another interesting dynamic starting from *I*_0_ = 1. *I*_*t*_ becomes larger than *Î* (overshoots) and then converges monotonically down to *Î*; in each case 0 < *df*(*I*)/*dt*|_*I*=*Î*_ < 1. For the parameters in the seventh row, there is oscillatory convergence to *Î* from *I*_0_ = 1 (−1 < *df*(*I*)/*dI*|_*I*=*Î*_ < 0), while in the last row there is straightforward monotone convergence to *Î*.

A continuous-time analog of the discrete-time recursion (11), in the form of a differential equation, substitutes *dI/dt* for *I*_*t*+1_ − *I*_*t*_ in (11). We then solve the resulting delay differential equation numerically using the VODE differential equation integrator in SciPy [36, 37] (source code available at https://github.com/yoavram/SanJose). Using the parameters in Table 2, Figures 1–4 compare the effect of the parameters on the trajectories of the discrete-time and continuous-time SIS model specified in (11). The number of time steps used in the computations illustrated in these figures is less than 250 in each case. In Figure 2 the delay is Δ = 2 while Figures 3 and 4 have delay Δ = 3. In the supplementary material, the discrete-time and continuous-time recursions of the SIR model are compared for short (Figures S1S–S4S) and much longer (Figures S1L–S4L) durations.

**Fig 1.**
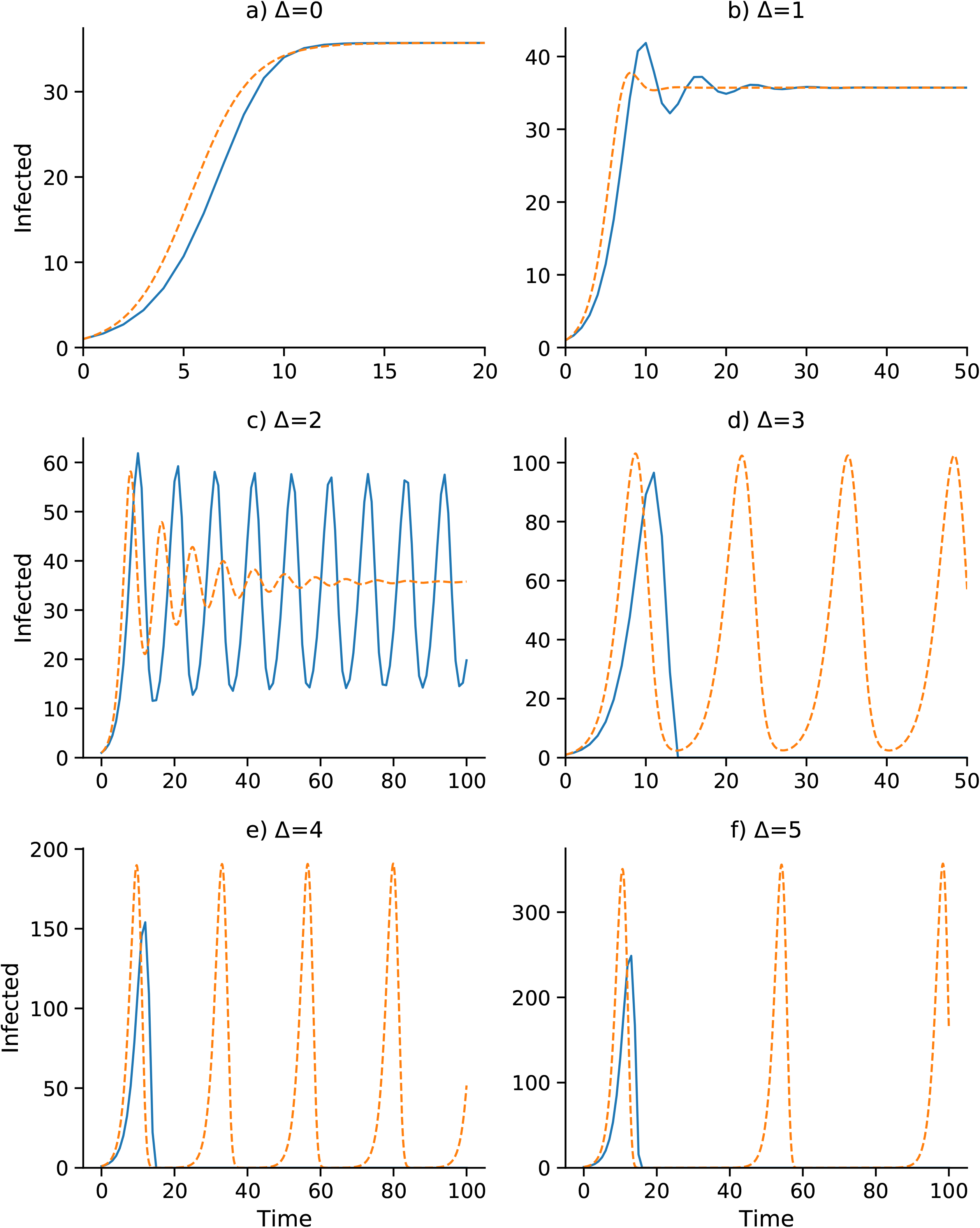
**Discrete-time SIS (blue) and continuous-time SIS (orange) dynamics** for delays Δ = 0 to Δ = 5 with *N* = 10, 000, *b*_0_ = 0.05, *γ* = 0.08, *ĉ* = 0.0015, *α* = 0.375, and *I*_0_ = 1. Here the epidemic equilibrium is *Î* = 35.72.

**Fig 2.**
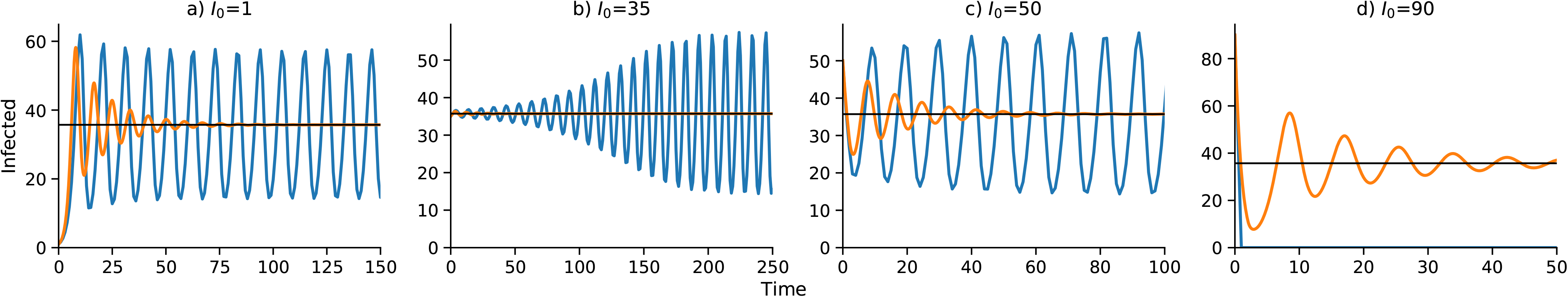
Effect of initial number of infecteds *I*_0_ on the dynamics for delay Δ = 2. Discrete- and continuous-time results are in blue and orange, respectively. Other parameters as in Figure 1. As in Figure 1, *Î* = 35.72.

**Fig 3.**
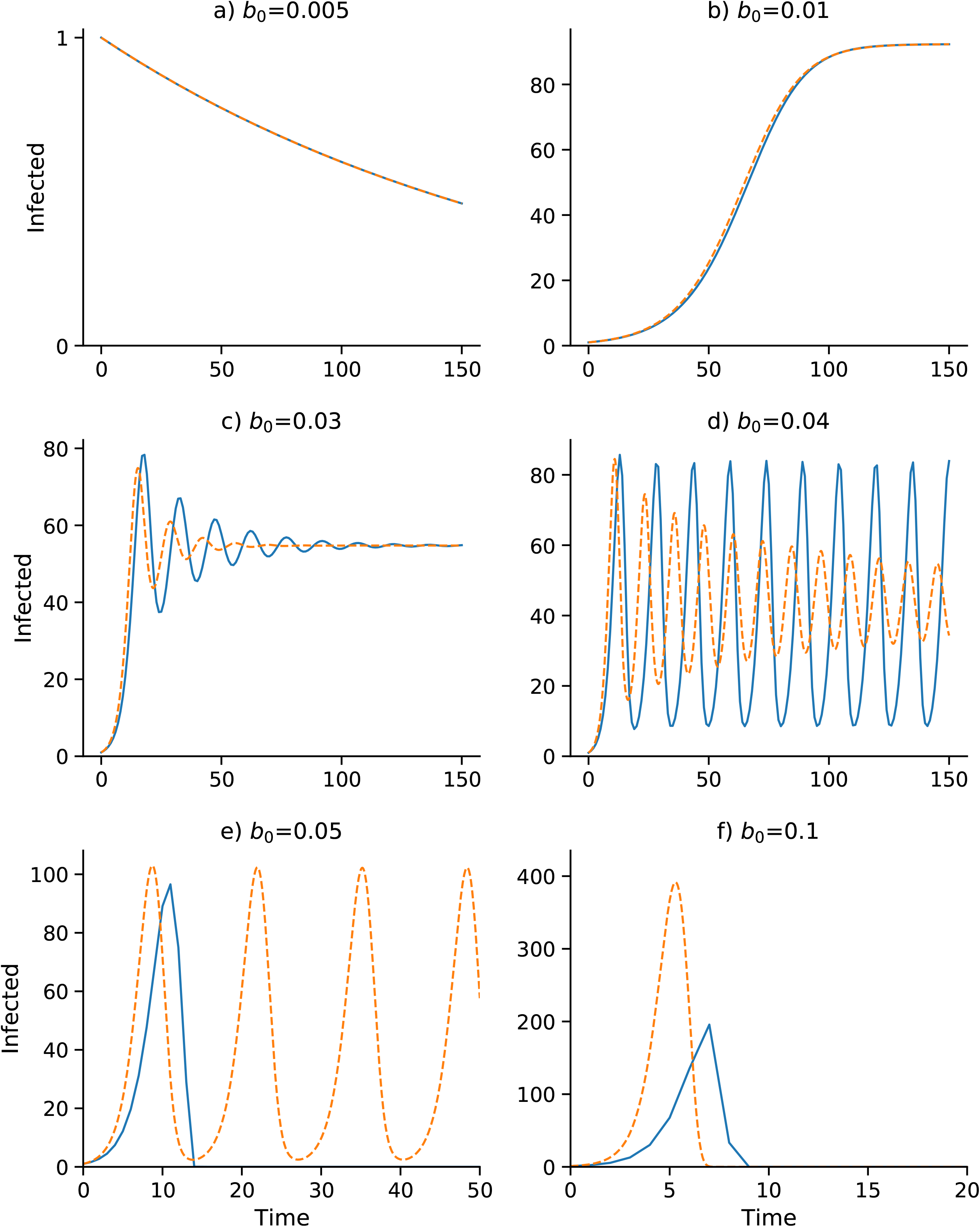
Effect of baseline contact rate *b*_0_ on dynamics with delay Δ = 3. Other parameters as in Figure 1 with *I*_0_ = 1. Discrete- and continuous-time results are in blue and orange, respectively. Note that *α* changes with *b*_0_ as *α* = *b*_0_*α*_2_/2*α*_1_: (a) *α* = 0.0375; (b) *α* = 0.075; (c) *α* = 0.225; (d) *α* = 0.3; (e) *α* = 0.375; (f) *α* = 0.75.

**Fig 4.**
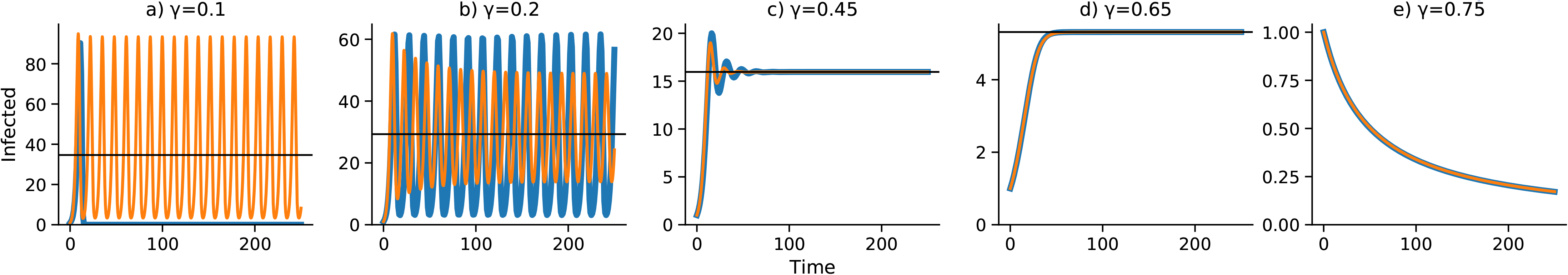
Effect of removal rate *γ* on dynamics with delay Δ = 3. Discrete- and continuous-time results are in blue and orange, respectively. Other parameters as in Figure 1 with *I*_0_ = 1.

In Figure 1, with no delay (Δ = 0) and a one-unit delay (Δ = 1), the discrete and continuous dynamics are very similar, both converging to *Î*. However, with Δ = 2 the differential equation oscillates into *Î* while the discrete-time recursion enters a regime of inexact cycling around *Î*, which appears to be a state of chaos. For Δ = 3 and Δ = 4, the discrete recursion “collapses”. In other words, *I*_*t*_ becomes negative and appears to go off to −∞; in Figure 1, this is cut off at *I* = 0. The continuous version, however, in these cases enters a stable cycle around *Î*. It is important to note that in Figure 1 for each panel the initial frequency was *I*_0_ = 1 infected individual. For Δ = 2, for example, with an initial value of *I*_0_ higher than about 73, instead of the inexact cycle, which is approached for smaller values of *I*_0_, the discrete recursion goes off and becomes negatively unbounded. This dependence of the dynamics on *I*_0_ is illustrated for Δ = 2 in Figure 2, where the continuous-time version of the SIS model (11) oscillates into *Î*. Two expanded views of the inexact cycling seen for *I*_0_ = 1 in Figure 2 are presented in Supplementary Figure S5.

Figures 3 and 4 focus on a delay of Δ = 3 and show the dependence of the discrete- and continuous-time dynamics on parameters *b*_0_ and *γ*, respectively. For *b*_0_ increasing from 0.005 to 0.05 the pattern of trajectories from *I*_0_ = 1 is remarkably similar to that for *γ* decreasing from 0.75 to 0.1. First, both converge to *Ĩ* = 0, then both converge to *Î*, then there is stable oscillation into *Î*. For *b*_0_ = 0.04 and *γ* = 0.2, however, the continuous trajectory enters a stable cycle while the discrete trajectory cycles inexactly around *Î*. For higher values of *I*_0_, however, the discrete-time trajectory may become unbounded. Finally, for *b*_0_ = 0.05 and *γ* = 0.75, the discrete-time trajectory goes of to −∞, but is shown stopped at 0, while the continuous case develops a stable cycle.

The discrete-time and continuous time trajectories for the SIR model (13–15) were studied with the same parameters as used in Figs. 1–4. Each computation is presented twice: first, for the same length of time as the SIS discrete- and continuous-time in Figs. 1–4, and second, for up to 5,000 time units. Graphs of the trajectories are shown in the Supplementary material, where S1S, S2S, S3S, and S4S refer to the shorter run times and S1L, S2L, S3L, and S4L refer to the longer run times. For the longer run times, as expected, both discrete-time and continuous-time versions of the SIR model eventually converge to *R* = *N* with no infecteds. Comparing the short-run figures with the long-run figures, the former are not good predictors of the latter in the SIR setting. The short-run behavior of the discrete-time model usually involves a great deal of cycling, which is difficult to see on the longer time scales. Supplementary Figs. S5S and S5L describe the SIR dynamics for the SIS version in Fig. 2, panel (a), with *I*_0_ = 1. Figs. S5S and S5L show the SIR versions of this figure run for a short and long time, respectively. In Fig. S5S there appears to be convergence to *Î*, but in Fig. S5L after about 500 time units, in both discrete- and continuous-time SIR versions, the number of infected begins to decline towards zero.

It is worth noting that if the total population size of *N* decreases over time, for example, if we take *N*(*t*) = *N*exp(−*zt*), with *z* = 50*b*_0_*ĉγ*, then the short-term dynamics of the SIS model in (11) begins to closely resemble the SIR version. This is illustrated in Supplementary Fig. S5N, where *b*_0_*, *ĉ*, γ* are, as in Figs. S5S and S5L, the same as in Fig. 2, panel (a). With *N* decreasing to zero, both *S* and *I* will approach zero in the SIS model, which explain its apparent similarity to the SIR model, where *S* and *I* approach zero as *R* approaches *N*.

## Discussion

This simple model of adaptive social contact produces two possible equilibria, one where there are zero infecteds, and the disease is eradicated, and a non-zero one between zero and *N*, the population size. These equilibria are stable under different conditions. Further, dynamics produced by this model are complex and subject to regime shifts across thresholds in the initial conditions and parameter settings. These dynamics include damped oscillation to the equilibrium, periodic oscillation, chaotic oscillation, and regression to positive or negative infinity.

Our model makes a number of simplifying assumptions. We assume, for example, that all individuals in the population will respond in the same fashion to government policy. We assume that governments choose a uniform contact rate according to an optimized utility function, which is homogeneous across all individuals in the population. Finally, we assume that the utility function is symmetric around the optimal number of contacts so that increasing or decreasing contacts above or below the target contact rate, respectively, yield the same reduction in utility. These assumptions allowed us to create the simplest possible model that includes adaptive behavior and time delay. In Holling’s heuristic distinction in ecology between tactical models, models built to be parameterized and predictive, and strategic models, which aim to be as simple as possible to highlight phenomenological generalities, this is a strategic model [38].

We note that the five distinct kinds of dynamical trajectories seen in these computational experiments come from a purely deterministic recursion. This means that oscillations and even erratic, near-chaotic dynamics and collapse in an epidemic may not necessarily be due to seasonality, complex agent-based interactions, changing or stochastic parameter values, demographic change, host immunity, or socio-cultural idiosyncracies. This dynamical behavior in number of infecteds can result from mathematical properties of a simple deterministic system with homogeneous endogenous behavior-change, similar to complex population dynamics of biological organisms [39]. The mathematical consistency with population dynamics suggests a parallel in ecology, that the indifference point for human behavior functions in a similar way to a carrying capacity in ecology, below which a population will tend to grow and above which a population will tend to shrink. For example, the Ricker Equation [40], commonly used in population dynamics to describe the growth of fish populations, exhibits similar complex dynamics and qualitative state thresholds. The existence of a non-zero, locally stable equilibrium in our model is consistent with economic epidemiology theory: if individuals are incentivized to change their behavior to protect themselves, they will, and they will cease to do this when they are not [10]. Further, our results show certain parameter sets can lead to limit-cycle dynamics, consistent with other negative feedback mechanisms with time delays [41, 42]. This is because the system is reacting to conditions that were true in the past, but not necessarily true in the present. In our discrete-time model, there is the added complexity that the non-zero equilibrium may be locally stable but not attained from a wide range of initial conditions, including the most natural one, namely a single infected individual.

Observed epidemic curves of many transient disease outbreaks typically inflect and go extinct, as opposed to this model that may oscillate perpetually or converge monotonically or cyclically to an endemic disease equilibrium. Our model is meant to demonstrate what effect incentives can have on infectious disease dynamics. Including institutional and public efforts that are incentivized to eradicate, rather than to optimize utility trade-offs, would alter the dynamics to look more like real-world epidemic curves. This may have a useful implication for policy. For example, beyond infectious diseases that remain endemic to society, outbreaks may also flare up once or multiple times, such as the double-peaked outbreaks of SARS in three countries in 2003 [43], and surges in fluctuations in COVID-19 cases globally [1]. There may be many causes for such double-peaked outbreaks, one of which may be a lapse in behavior-change after the epidemic begins to die down due to decreasing incentives [16], as represented in our simple theoretical model. This is consistent with findings that voluntary vaccination programs suffer from decreasing incentives to participate as prevalence decreases [44, 45]. It should be noted that the continuous-time version of our model can support a stable cyclic epidemic whose interpretation in empirical terms will depend on the time scale, and hence on the meaning of the delay, Δ.

One of the responsibilities of infectious disease modelers (e.g. COVID-19 modelers) is to predict and project forward what epidemics will do in the future in order to better assist in the proper and strategic allocation of preventative resources. COVID-19 models have often proved wrong by orders of magnitude because they lack the means to account for adaptive response. An insight from this model, however, is that prediction becomes very difficult, perhaps impossible, if we allow for adaptive behavior-change because the system is qualitatively sensitive to small differences in values of key parameters. These parameters are very hard to measure precisely; they change depending on the disease system and context and their inference is generally subject to large errors. Further, we don’t know how policy-makers weight the economic trade-offs against the public health priorities (i.e., the ratio between *α*_1_ and *α*_2_ in our model) to arrive at new policy recommendations.

To maximize the ability to predict and minimize loss of life or morbidity, outbreak response should not only seek to minimize the reproduction number, but also the length of time taken to gather and distribute information. Another approach would be to use a predetermined strategy for the contact rate, as opposed to a contact rate that depends on the number of infecteds.

In our model, complex dynamic regimes occur more often when there is a time delay. If behavior-change arises from fear and fear is triggered by high local mortality and high local prevalence, such delays seem plausible since death and incubation periods are lagging epidemiological indicators. Lags mean that people can respond sluggishly to an unfolding epidemic crisis, but they also mean that people can abandon protective behaviors prematurely. Developing approaches to incentivize protective behavior throughout the duration of any lag introduced by the natural history of the infection (or otherwise) should be a priority in applied research. This paper represents a first step in understanding endogenous behavior-change and time-lagged protective behavior, and we anticipate further developments along these lines that could incorporate long incubation periods and/or recognition of asymptomatic transmission.

## Supporting information

**S1S Fig. Discrete-time (blue) and continuous-time (orange) versions of the SIR model Eqs. (13)–(15).** Parameters are the same as in Fig. 1.

**S1L Fig. Longer time runs for the SIR cases in Figure S1S.** The discrete-time (blue) and continuous-time (orange) trajectories are very similar for Δ = ‘, 1, 2. For Δ = 3, 4, 5, the discrete-time trajectories are stopped at *I* = 0, as they go off to −∞. The continuous-time cases all converge to zero infecteds.

**S2S Fig. SIR version of the SIS model in Fig. 2 with Δ = 2.** Discrete-time (blue) and continuous-time (orange) trajectories are similar to the SIS graphs. Parameters as in Fig. 2.

**S2L Fig. SIR version of the SIS model in Fig. 2 with Δ = 2 run for 5,000 time units.** Both discrete-time (blue) and continuous-time (orange) graphs converge to zero infecteds. Parameters as in Fig. 2.

**S3S Fig. SIR version of the SIS model in Fig. 3.** Discrete-time (blue) and continuous-time (orange) trajectories are similar to the SIS graphs in Fig. 3. Parameters as in Fig. 3.

**S3L Fig. SIR models in Figure S3S run for 5,000 time units.** Assumptions as in Fig. S3S.

**S4S Fig. Effect of removal rate *γ* on discrete-time (blue) and continuous-time (orange) versions of the SIR model.** Note the compression of the cycles seen in Figure 4 and the earlier decline to zero infecteds.

**S4L Fig. SIR models in Figure S4S run for 4,000 generations.** All runs eventually approach zero infecteds. Parameters as in Figure 4.

**S5 Fig. Dynamics with delay Δ = 2 and initial number of infecteds *I*_0_ = 1 in the SIS model (same as Figure 2, panel (a)).** (**a**): Return map showing more than one *I*_*t*+1_ value for each value of *I*_*t*_. (**b**): Comparing the “elliptical” dynamics in part (a) with continuous-time damped oscillation (orange) to equilibrium *Î* = 35.72. Other parameters as in Figure 2. This figure is the same as Figure 2, panel (a).

**S5S Fig. SIR versions of discrete-time (blue) and continuous-time (orange) versions of the SIS model in Figure 2, panel (a).** Note the apparent approach to *Î*.

**S5L Fig. Same as Figure S5S run for 5,000 time units.** Both discrete-time (blue) and continuous-time (orange) trajectories eventually approach *R* = *N* .

**S5N Fig. SIS model (recursion (11)) with *N* decreasing over time.** This uses the same parameters as in Figure 2 but sets *N* = *N*(*t*) = exp(−*zt*), where *z* = 50*γb*_0_*ĉ* with *γ* = 0.08, *b*_0_ = 0.05, *ĉ* = 0.0015. Note the similarity to Figure S5S.

## S1 Appendix. Local stability of *Î*

Recall the recursion (11):

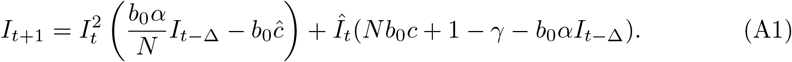

In the neighborhood of the equilibrium *Î*, write *I*_*t*_ = *Î* + *ε*_*t*_ and *I*_*t*−Δ_ = *Î* + *ε*_*t*−Δ_, where *ε*_*t*_ and *ε*_*t*−Δ_ are small enough that quadratic terms in them can be neglected in the expression for *I*_*t*+1_ = *Î* + *ε*_*t*+1_. The linear approximation to (A1) is then

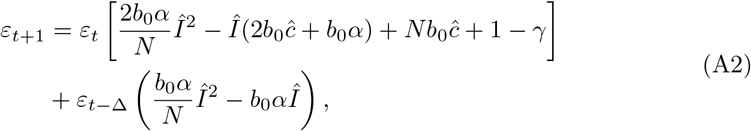

and in the case Δ = 0, this reduces to

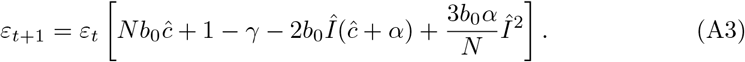

We focus first on Δ = 0 and write (A3) as *ε*_*t*+1_ = *ε*_*t*_*L*(*Î*). Recall that *Î* satisfies Eq. (17), and substituting *γ* from (17) into *L*(*Î*), we obtain

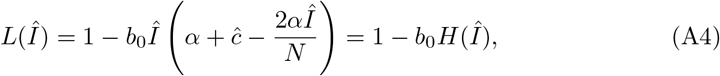

where

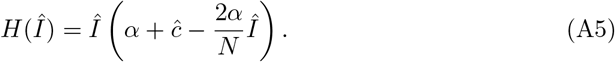

Clearly *N* ≥ *Î*, and since *c*\* must be positive, *ĉ* > *α*Î**/*N* . Hence *H*(*Î*) > 0 and, for local stability of *Î*, the remaining condition for |*L*(*Î*)| < 1 is *b*_0_*H*(*Î*) < 2. Direct substitution of *Î* gives *b*_0_*H*(*Î*) < 2 if

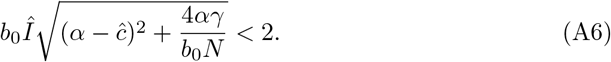

Now we turn to the general case Δ ≠ 0 and Eq. (A2), which we write as

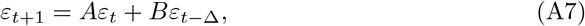

where *A* and *B* are the corresponding terms on the right side of (A2). Eq. (A7) is a homogeneous linear recursion, since, given *Î* and all the parameters, *A* and *B* are constants with respect to time. Local stability of *Î* is then determined by the properties of recursion (A7), whose solution first involves solving its characteristic equation

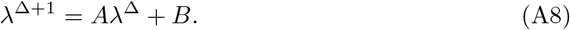

In principle there are Δ + 1 real or complex roots of (A8), which we represent as *λ*_1_, *λ*_2_, …, *λ*_Δ+1_, and the solution of (A7) can be written as

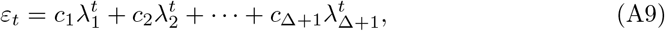

where *c*_*i*_ are found from the initial conditions. Convergence to, and hence local stability of *Î*, is determined by the magnitude of the absolute value (if real) or modulus (if complex) of the roots *λ*_1_, *λ*_2_, … , *λ*_Δ+1_: *Î* is locally stable if the largest among the Δ + 1 of these is less than unity.

In Table 2, results of numerically iterating the complete recursion (11) are listed for the delay Δ varying from Δ = 0 to Δ = 4, all starting from *I*_0_ = 1, with *N* = 10, 000 and the stated parameters. Figure 3 illustrates the discrete- and continuous-time dynamics summarized in Table 2. With these values, *Î* = 35.7180 and we obtain *A* = 0.9997 and *B* = −0.6673. Then, for Δ = 0, Eq. (A7) gives *ε*_*t*_ = 0.3324*ε*_*t*−1_, which entails that convergence to *Î* is locally monotone. With Δ = 1, the characteristic polynomial is a quadratic,

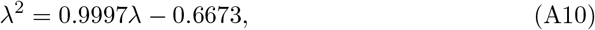

with complex roots 0.4999 ± 0.6461*i* whose modulus is 0.8169, which is less than 1. The complexity implies cyclic behavior, and since the modulus is less than one, we see locally damped oscillatory convergence to *Î*.

For Δ = 2, the characteristic equation is the cubic

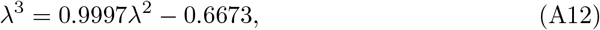

which has one real root 0.6383 and complex roots 0.8190 ± 0.6122*i*. Here the modulus of the complex roots is 1.0225, which is greater than unity so that *Î* is not locally stable. In this case the dynamics depend on the initial value *I*_0_. If *I*_0_ < 72, *I*_*t*_ oscillates but not in a stable cycle. If *I*_0_ > 73, the oscillation becomes unbounded.

When Δ = 3, the four roots of the characteristic polynomial

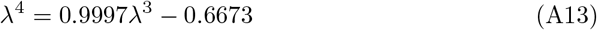

are all complex: −0.4566 ± 0.5966*i* and 0.9564 ± 0.5173*i*. The modulus of the second pair of complex roots is greater than 1. For Δ = 4, the five roots of

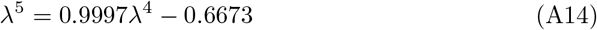

are −0.7823, −0.1301 ± 0.8212*i*, and 1.0211 ± 0.4376*i*. Here again the largest modulus is 1.1109, greater than 1. Thus for both Δ = 3 and 4, *Î* cannot be locally stable, and for these delay times the recursion can oscillate wildly becoming negatively and positively unbounded for some starting values *I*_0_.

## Acknowledgments

This research was supported in part by the Morrison Institute for Population and Research Studies (MF), by Israel Science Foundation grants 552/19 and 3811/19 (YR), and by a Graduate Research Fellowship from the National Science Foundation (RA). The authors thank Kaleda Krebs Denton and W. Brian Arthur for helpful comments on an earlier draft of the manuscript.

